# Evolutionary implications of recombination differences across diverging populations of *Anopheles*

**DOI:** 10.1101/2021.02.04.429659

**Authors:** Joel T. Nelson, Omar E. Cornejo, Ag1000G Consortium

## Abstract

Recombination is one of the main evolutionary mechanisms responsible for changing the genomic architecture of populations; and in essence, it is the main mechanism by which novel combinations of alleles, haplotypes, are formed. A clear picture that has emerged across study systems is that recombination is highly variable, even among closely related species. However, it is only until very recently that we have started to understand how recombination variation between populations of the same species impact genetic diversity and divergence. Here, we used whole-genome sequence data to build fine-scale recombination maps for nine populations within two species of *Anopheles*, *Anopheles gambiae* and *Anopheles coluzzii*. The genome-wide recombination averages were on the same order of magnitude for all populations except one. Yet, we identified significant differences in fine-scale recombination rates among all population comparisons. We report that effective population sizes, and presence of a chromosomal inversion has major contribution to recombination rate variation along the genome and across populations. We identified over 400 highly variable recombination hotspots across all populations, where only 9.6% are shared between two or more populations. Additionally, our results are consistent with recombination hotspots contributing to both genetic diversity and absolute divergence (dxy) between populations and species of *Anopheles*. However, we also show that recombination has a small impact on population genetic differentiation as estimated with F_ST_. The minimal impact that recombination has on genetic differentiation across populations represents the first empirical evidence against recent theoretical work suggesting that variation in recombination along the genome can mask or impair our ability to detect signatures of selection. Our findings add new understanding to how recombination rates vary within species, and how this major evolutionary mechanism can maintain and contribute to genetic variation and divergence within a prominent malaria vector.

## Introduction

Meiotic recombination is the main evolutionary mechanism that shapes haplotypic variation in sexually reproducing species (Felsenstein 1974; San Filippo et al. 2008; Beeson et al. 2019). A main result of recombination is the movement of alleles onto different genetic backgrounds, which has the potential to create novel combinations of beneficial haplotypes and novel haplotypes (Posada et al. 2002; Smagulova et al. 2016; Korunes and Noor 2017). Recombination can reduce interference between linked loci, thereof reducing mutational load, and it allows for beneficial mutations to sweep through a population more rapidly (Gabriel et al. 1993; McVean, G. A. and Charlesworth 2000; Otto and Barton 2001; Hartfield and Keightley 2012; Ritz et al. 2017). It has been suggested that elevated recombination rates can contribute to increased patterns of genetic diversity and divergence, though this relationship is not well understood (Smukowski and Noor 2011). Across most species, recombination is highly variable within and across chromosomes but tends to cluster in local regions identified as recombination hotspots (Hellsten et al. 2013; Beeson et al. 2019). Both fine-scale recombination rates and hotspots are poorly conserved across mammals and plants with significant differences even among closely related species (Spencer et al. 2006; Choi et al. 2016; Beeson et al. 2019; Dreissig et al. 2019). However, less is known about the onset of variation in recombination rates across diverging populations of the same species (Schwarzkopf et al. 2020) and how it impacts patterns of genetic variation. Comparing recombination maps across multiple populations of humans has recently been investigated; however, this question has not been applied to other model systems and was aimed at identifying the impact of demographic changes on the recombination landscape and not how elevated recombination impacts patterns of diversity and divergence (Spencer et al. 2006). Thus, comparing recombination maps across closely related populations and quantifying its impact on genetic variation is imperative for elucidating how this mechanism shapes patterns of evolution.

Comparing recombination maps across closely related populations could help predict shifts in the recombination landscape that are responsible for species-level differences in recombination (Smukowski and Noor 2011). For example, comparing the recombination landscape across genomes with varying selective sweeps, effective populations sizes, and chromosomal inversions. These aspects of genomic evolution are known to impact recombination events, however the contribution they have towards recombination rate differentiation across populations in generally unknown (Keightley and Otto 2006; Feder and Nosil 2009; Yang et al. 2018). Furthermore, comparing recombination landscapes among populations can also have significant contributions towards understanding the impact that elevated recombination rates have on patterns of genetic diversity and divergence (Brown and Jiricny 1987; Brown et al. 1989; Papavasiliou and Schatz 2000; Smukowski and Noor 2011). For instance, because of the hitchhiking effects of linked selection, local regions of the genome with significantly larger recombination rates (recombination hotspots) may act as reservoirs for genetic diversity. Previous studies have identified an increase in nucleotide diversity within regions of elevated recombination; however, there is a discord among studies for the number of differences between species within the same regions (Smukowski and Noor 2011; Roesti et al. 2013). In concordance to genetic diversity and divergence, little is known about the impact that elevated recombination rates have on localized patterns of genetic differentiation (F_ST_); and if lower levels of differentiation are expected in regions of the genome with elevated recombination rates due to reduced linkage. Thus, understanding these patterns is an important step in elucidating the early onset of recombination rate evolution among closely related populations and the impact that elevated rates have on levels of genetic diversity and divergence. A perhaps more subtle, but equally important relevance of recombination in modern genomic studies is the potential impact that differences in recombination have on our ability to identify signatures of hard or soft sweeps in the genome. Using simulations, it has been shown that recombination acting on regions close to a hard sweep can generate patterns of polymorphism that resemble those found in soft sweeps, rendering them indistinguishable (Pennings and Hermisson 2006; Schrider et al. 2015).

To address recombination rate variation among closely related populations we used sequence data from two different species of *Anopheles*, *A. gambiae* and *A. coluzzii;* which are the major contributors to the spread of malaria (Holt et al. 2002; Anopheles gambiae 1000 Genomes Consortium 2017). There are several known populations of *A. gambiae* and *A. coluzzii* that have a large distribution throughout Africa, spanning across prominent ecological gradients (Anopheles gambiae 1000 Genomes Consortium 2017). Previous studies have identified several populations that have undergone expansions and bottlenecks, as well as containing the presence of selective sweeps and chromosomal inversions, characteristics that contribute to differences in the recombination architecture of populations (Lehmann et al. 1999; Anopheles gambiae 1000 Genomes Consortium 2017). Populations within this system have low background levels of genetic differentiation (White et al. 2010; Anopheles gambiae 1000 Genomes Consortium 2017). The combination of multiple population pairwise comparisons and low levels of genetic differentiation allows for robust detection of increased genetic differentiation and divergence in localized regions of the genome where recombination is elevated (e.g. recombination hotspots). There is a large body of evidence showing a correlation between recombination and nucleotide diversity in *Drosophila melanogaster* (Begun and Aquadro 1992; Andolfatto and Przeworski 2001), with no consequence on divergence, which has led to suggest that natural selection is an important force structuring patterns of polymorphism along the genome (Wright et al. 2006). The pattern described for *Drosophila* has not always been replicated in other organisms, where divergence is at times correlated with recombination, suggesting a lack of generality on this observation (Nachman et al. 1998; Nachman 2001; Lercher and Hurst 2002; Cutter and Payseur 2003; Hellmann, Ines et al. 2003; Hellmann, I. et al. 2005). We believe that a more extensive analysis of the architecture of recombination and genome-wide polymorphism is necessary to shed light into this phenomenon.

Here, we use whole genome data to infer fine-scale recombination maps across nine diverging populations of *A. gambiae* and *A. coluzzii* (Figure 1). Specifically, we 1) quantified the recombination landscape across populations and identified the impact that different evolutionary/genomic features have on recombination rates, 2) identified regions of the genome where recombination rates were significantly greater than average background levels (hotspots), and 3) quantified the impact of elevated recombination on population diversity and divergence. We expected to see large variations in recombination rates, even among closely related populations (fine-scale), yet smaller difference when comparing genome-wide rates (broad-scale). Based on what has been observed in studies in other organisms, we expected that differences in recombination hotspots will be similar to variation in recombination rates where most hotspots would be unique to a specific population. Simulation studies have shown that recombination differences can reduce our ability to detect regions of the genome under selection, evidenced by excessive differentiation. We expect that our analyses of recombination will allow to investigate the extent of these expectations in natural populations.

**Figure 1.**
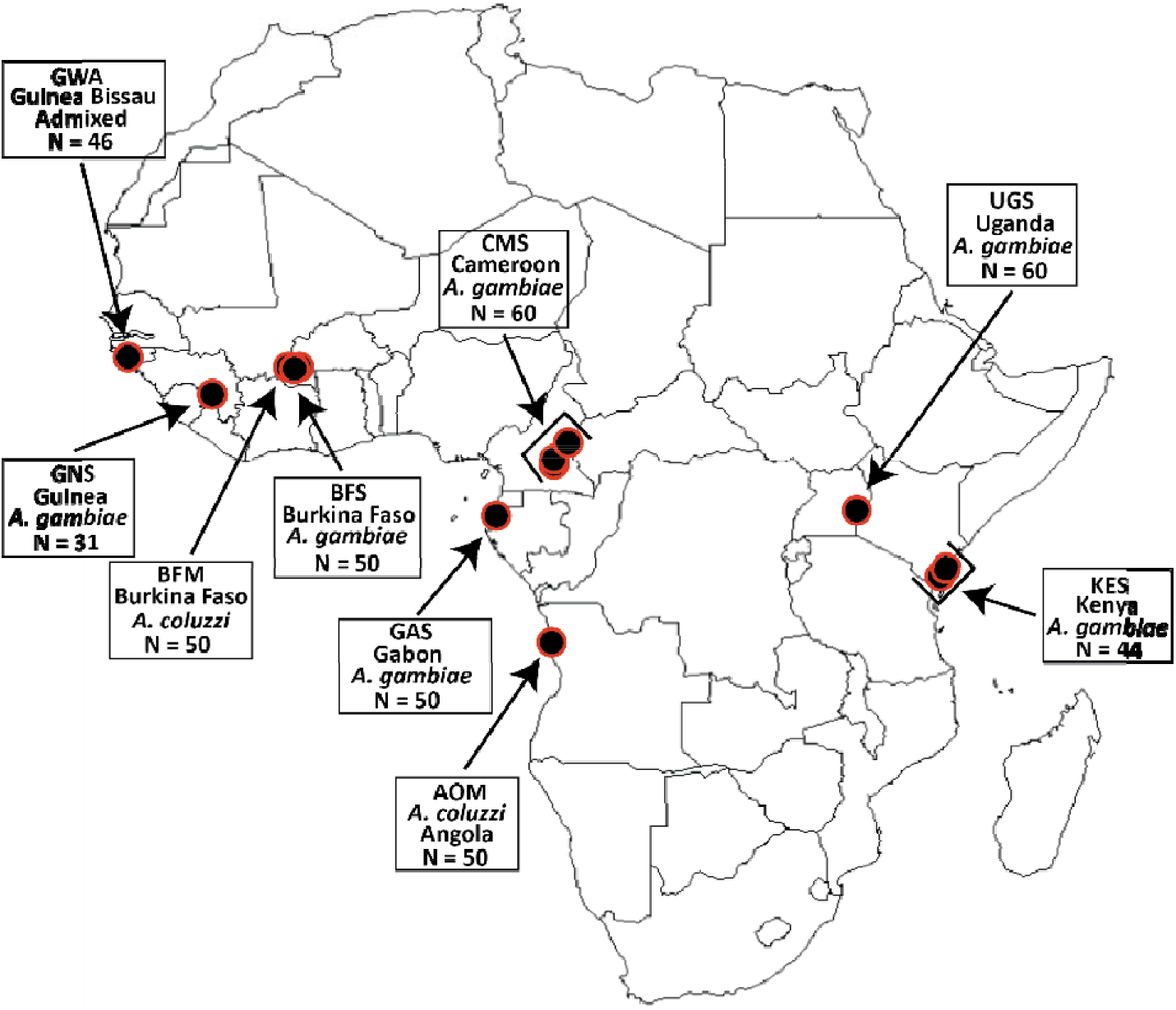
Map of Africa depicting the locations and sample sizes of each populations used for estimating recombination rates. Note that for several populations there multiple sampling locations.

## Results

### Comparing fine-scale recombination maps across diverging populations

Average population-scaled recombination rate (ρ = *4Ner*, where *Ne* is the effective population size and *r* is the per site recombination rate; measures one recombination event per kb) across the entire genome was 70.9 ρ. Population averages ranged from 0.58 ρ (KES) to 140.26 ρ (CMS) (Table 1 & 2) (see Supplementary Figure S1 – S3 for summarized recombination maps for each species). Genome-wide recombination distributions followed a positively skewed distributions where most recombination events throughout the genome were low with the exceptions to localized regions with elevated recombination (Figure 2). Results from our Wilcoxon Rank test suggests significant difference in recombination rates across all population comparisons (p < 7.39e-14) (Supplementary Table S1 & S2). After accounting for effective population size average per-site recombination rates (*r*) were all on the same order of magnitude expect for KES (Table 2). Average recombination rates (ρ/bp) were roughly two orders of magnitude lower than fine-scale estimates across species of *Drosophila* (Chan et al. 2012). Indeed, fine-scale recombination rates within *Anopheles* were in the lower quantile when including recombination estimates from other insect systems (Stapley et al. 2017).

**Table 1.** The average population-scaled (ρ) recombination rate for the entire genome, each autosomal chromosome arm, and the X chromosome.

| Population | $\rho$ (avg.) | 2L | 2R | 3L | 3R | X | Variance |
| --- | --- | --- | --- | --- | --- | --- | --- |
| **AOM** | 7.8 | 6.5 | 7.4 | 9.9 | 9.2 | 6.1 | 39.7765 |
| **BFM** | 111 | 107.1 | 60.7 | 138 | 131.9 | 117.5 | 6910.2 |
| **GWA** | 64.6 | 20.4 | 16.1 | 95.4 | 89 | 102.2 | 4965.31 |
| **BFS** | 119.8 | 70.5 | 107.8 | 151.2 | 145.3 | 124.1 | 10373.1 |
| **CMS** | 129.3 | 58.3 | 105.8 | 158.9 | 159.5 | 164 | 12530.1 |
| **GAS** | 4.4 | 4.4 | 4.5 | 5.3 | 5.6 | 2.1 | 14.2874 |
| **GNS** | 110.9 | 54.4 | 93.8 | 125.5 | 124 | 157 | 7618.57 |
| **KES** | 0.5 | 0.3 | 0.3 | 0.3 | 0.2 | 1.3 | 1.69677 |
| **UGS** | 89.9 | 36 | 85.6 | 78.6 | 113.3 | 135.8 | 8032.67 |
| **Average** | 70.9 | 39.8 | 53.6 | 84.8 | 86.5 | 90 | 5609.52 |

**Table 2.**
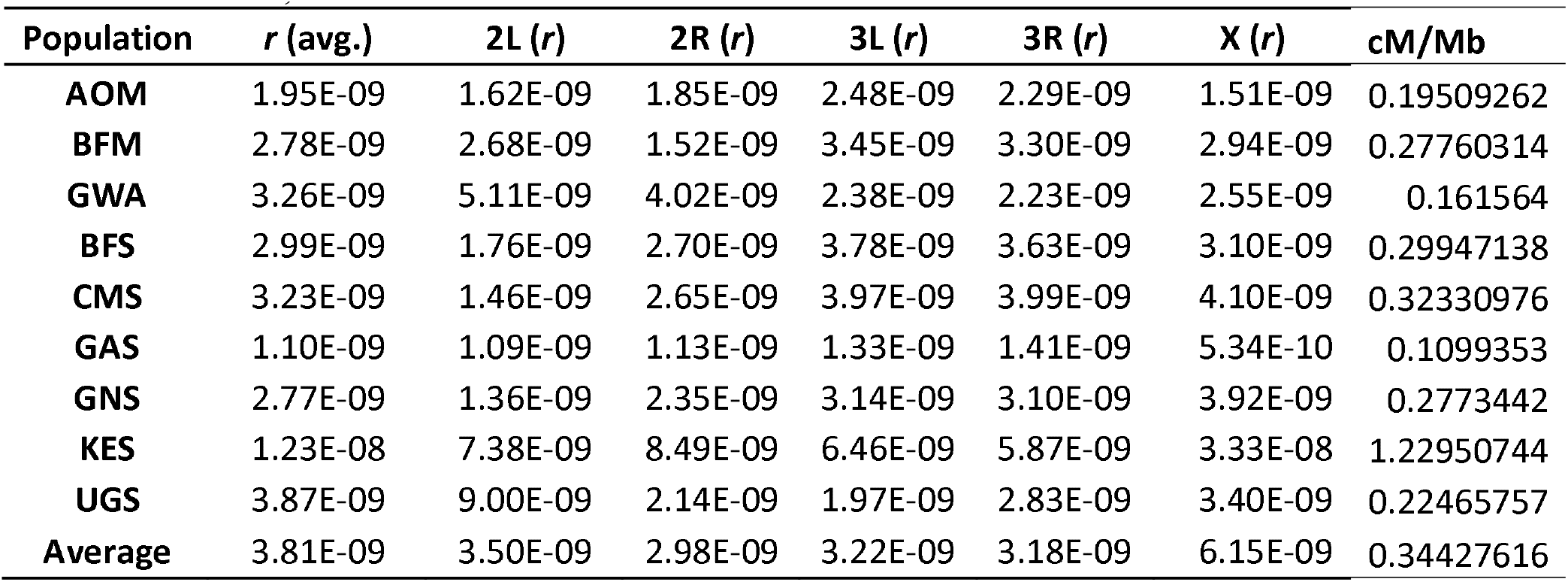
The average per-site (*r*) recombination rate for the entire genome, each autosomal chromosome arm, and the X chromosome.

| Population | $r$ (avg.) | 2L ($r$) | 2R ($r$) | 3L ($r$) | 3R ($r$) | X ($r$) | cM/Mb |
| --- | --- | --- | --- | --- | --- | --- | --- |
| **AOM** | 1.95E-09 | 1.62E-09 | 1.85E-09 | 2.48E-09 | 2.29E-09 | 1.51E-09 | 0.19509262 |
| **BFM** | 2.78E-09 | 2.68E-09 | 1.52E-09 | 3.45E-09 | 3.30E-09 | 2.94E-09 | 0.27760314 |
| **GWA** | 3.26E-09 | 5.11E-09 | 4.02E-09 | 2.38E-09 | 2.23E-09 | 2.55E-09 | 0.161564 |
| **BFS** | 2.99E-09 | 1.76E-09 | 2.70E-09 | 3.78E-09 | 3.63E-09 | 3.10E-09 | 0.29947138 |
| **CMS** | 3.23E-09 | 1.46E-09 | 2.65E-09 | 3.97E-09 | 3.99E-09 | 4.10E-09 | 0.32330976 |
| **GAS** | 1.10E-09 | 1.09E-09 | 1.13E-09 | 1.33E-09 | 1.41E-09 | 5.34E-10 | 0.1099353 |
| **GNS** | 2.77E-09 | 1.36E-09 | 2.35E-09 | 3.14E-09 | 3.10E-09 | 3.92E-09 | 0.2773442 |
| **KES** | 1.23E-08 | 7.38E-09 | 8.49E-09 | 6.46E-09 | 5.87E-09 | 3.33E-08 | 1.22950744 |
| **UGS** | 3.87E-09 | 9.00E-09 | 2.14E-09 | 1.97E-09 | 2.83E-09 | 3.40E-09 | 0.22465757 |
| **Average** | 3.81E-09 | 3.50E-09 | 2.98E-09 | 3.22E-09 | 3.18E-09 | 6.15E-09 | 0.34427616 |

**Figure 2.**
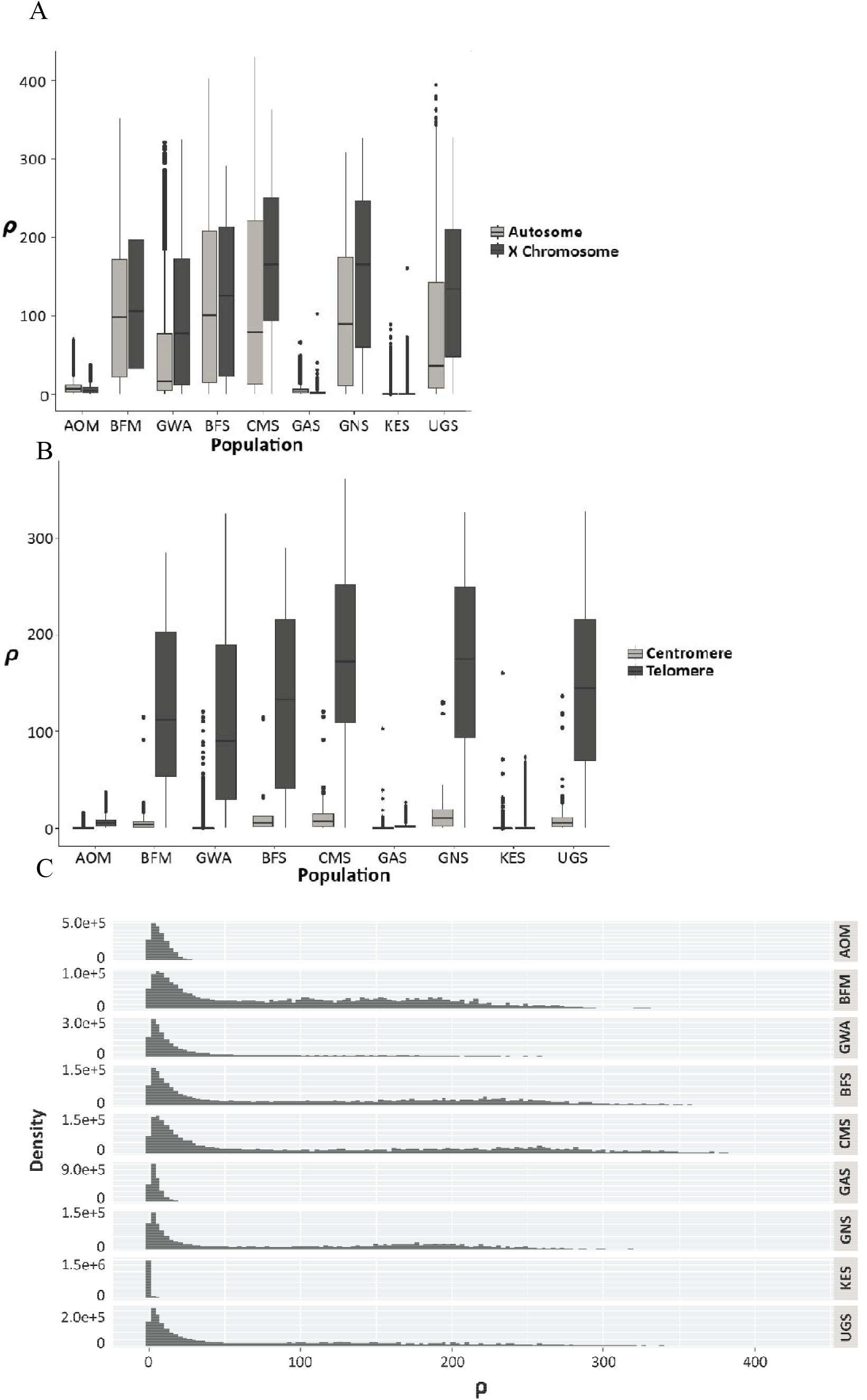
Top pannel (A) dipicts a boxplpot showing the degree of variation in population-scaled recombination rates within and across populations for the autosomal genome and the X chromosome. The middle pannel (B) shows the difference in population-scaled recombintion rates between the first 18Mb (Telomere) and the last 6Mb (Centromere) of the X chromosome for each population. The bottom pannel (C) represents individual histograms of the genome-wide (autosomal and the X chromosome) recombination rates for each population. Here, ρ is the population-scaled recombination rate which is the product of *4Ner*, where Ne is the effective population size and *r* is the per-site recombination rate.

### Ne, chromosomal location, inversions influence recombination variation, and sweeps

Populations with a smaller effective population size (AOM, GAS, and KES) exhibited a lower variance across the recombination landscape, while populations with a large effective population size reported nearly 100 times larger variance in the recombination landscape (Figure 2).When considering chromosomal location, we identified large reductions in recombination rates within centromeric regions of each chromosome, however, this patter was more prominent along the X chromosome (Figure 2 & 3). As expected, for populations that were polymorphic for chromosomal inversions, all recombination estimates (with the exception to BFS) fell below the genome average within inverted regions and then increased once outside of the inversion (Table 3; Figure 3). This pattern was not observed for populations that were homozygous for the inverted or wild-type genotypes (Figure 3), further confirming the characterization of the polymorphism of the inversion in these populations. Furthermore, Wilcoxon Rank test suggest significant difference among all pairwise comparisons in recombination rates within genomic regions containing the 2La and 2Rb inversion (p < 2.2e-16) (Supplementary Table S3 & S4). Lastly, we also showed that recombination rates were largely suppressed within insecticide resistant (IR) genes that have been identified as under strong selection in previous studies (Anopheles gambiae 1000 Genomes Consortium 2017). For example, the decrease in recombination rates in a 1 Mb region within the Cyp6p gene and a 3Mb region within the Cyp9k1 gene (Figure 4; also see Supplementary Figure S4-S9). For IR genes that were putatively evolving under neutral conditions (or weak signals of selection), we did not identify a prominent reduction in recombination rates or change in the topography of the recombination map.

**Table 3.** Averaged population-scaled recombination rate for the seven different inversions across smapled populations of *Anopheles*. The shaded boxes represents populations where there are at least three heterozygous genotypes for the 2Rb or the 2La inversion.

| Population | $\rho$<br>(avg.) | 2Rb(avg.) | 2La(avg.) |
| --- | --- | --- | --- |
| AOM | 8.22 | 11.17 | 7.06 |
| BFM | 104.54 | 20.41 | 103.77 |
| GWA | 50.61 | 15.58 | 8.29 |
| BFS | 117.71 | 146.71 | 32.79 |
| CMS | 118.96 | 87.55 | 15.04 |
| GAS | 4.95 | 5.45 | 4.22 |
| GNS | 98.78 | 73.21 | 7.84 |
| KES | 0.29 | 0.30 | 0.27 |
| UGS | 80.60 | 56.32 | 11.40 |

**Figure 3.**
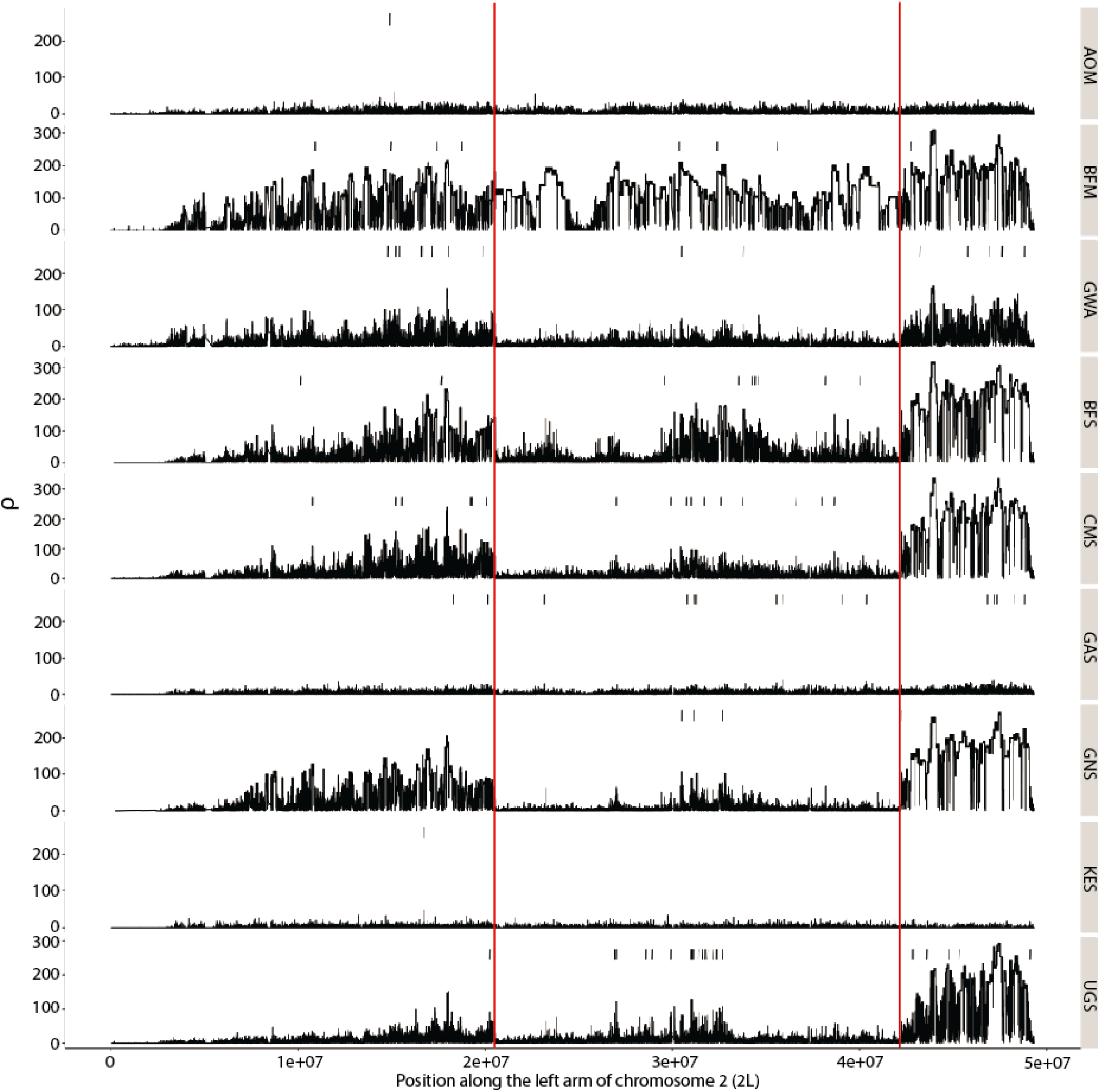
The estimated population-scaled recombination map of the left arm of the second chromsome for all populations. The small black vertical lines at the top of each map represents the location of a recombination hotspot identified from LDhot. The red lines denote the location of the 2La chromosomal inversions.

**Figure 4.**
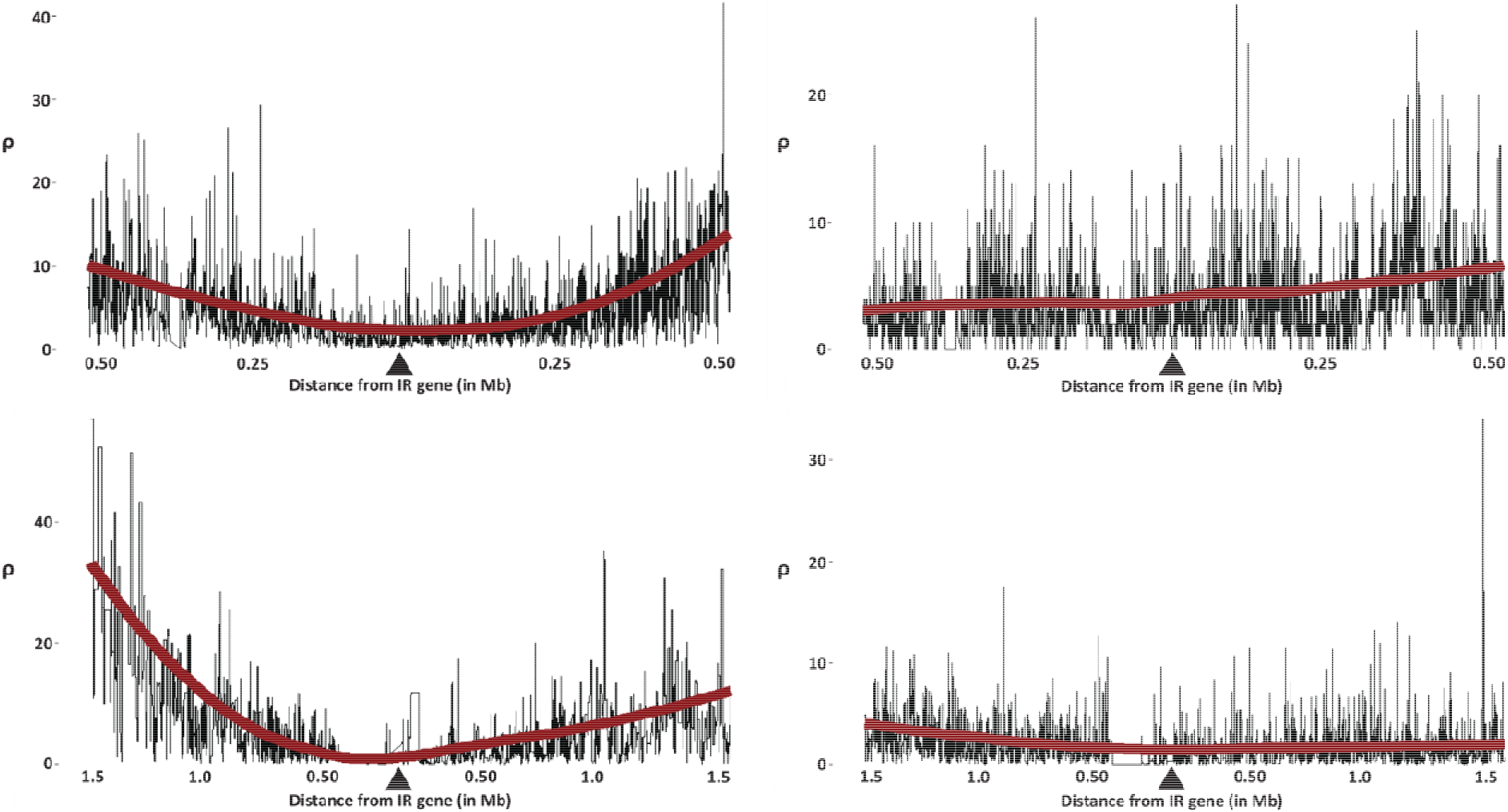
Population-scaled recombination rate for a genomic window containing two known IR genes that are under selection in some populations but are evolving neutrally in others. The red line denotes a smoothed average of recombination rates for the given region. Top panel shows the changes in recombination rate in a 1 Mb window containing the Cyp6p gene. Note that this gene shows evidence of selection in UGS (top left) but not in GAS (top right). The lower panel shows the changes in recombination rate in a 3 Mb window containing the Cyp9k1 gene. Previous work has found evidence for selection in BFM but weakly selected (or nearly neutral) in AOM. Here, the black triangle denotes the position of the given IR gene.

### Identifying unique and shared recombination hotspots

We identified a total of 435 robust signatures consistent with a recombination hotspot (Figure 5; Supplementary Table S5). In most populations, the average recombination rate within hotspots was at least twice as large as the adjacent background; however, the average recombination rates within hotspots were only slightly larger than the average genome-wide recombination rate across all populations (HS = ~71.0 ρ, genome = ~70.0 ρ). Moreover, for autosomal chromosomes, we identified 383 hotspots with an average of 48 hotspots per population. Of the 383 autosomal hotpots, 344 were unique to a single population while 39 were shared between two or more populations. For the X chromosome, we retained a total of 52 recombination hotspots with an average of six hotspots per population. Among the 52 hotspots, 51 were unique to a single population while only a single hotspot was shared between two populations (GWA and KES). A total of 26 population pairwise comparisons shared recombination hotpots.

**Figure 5.**
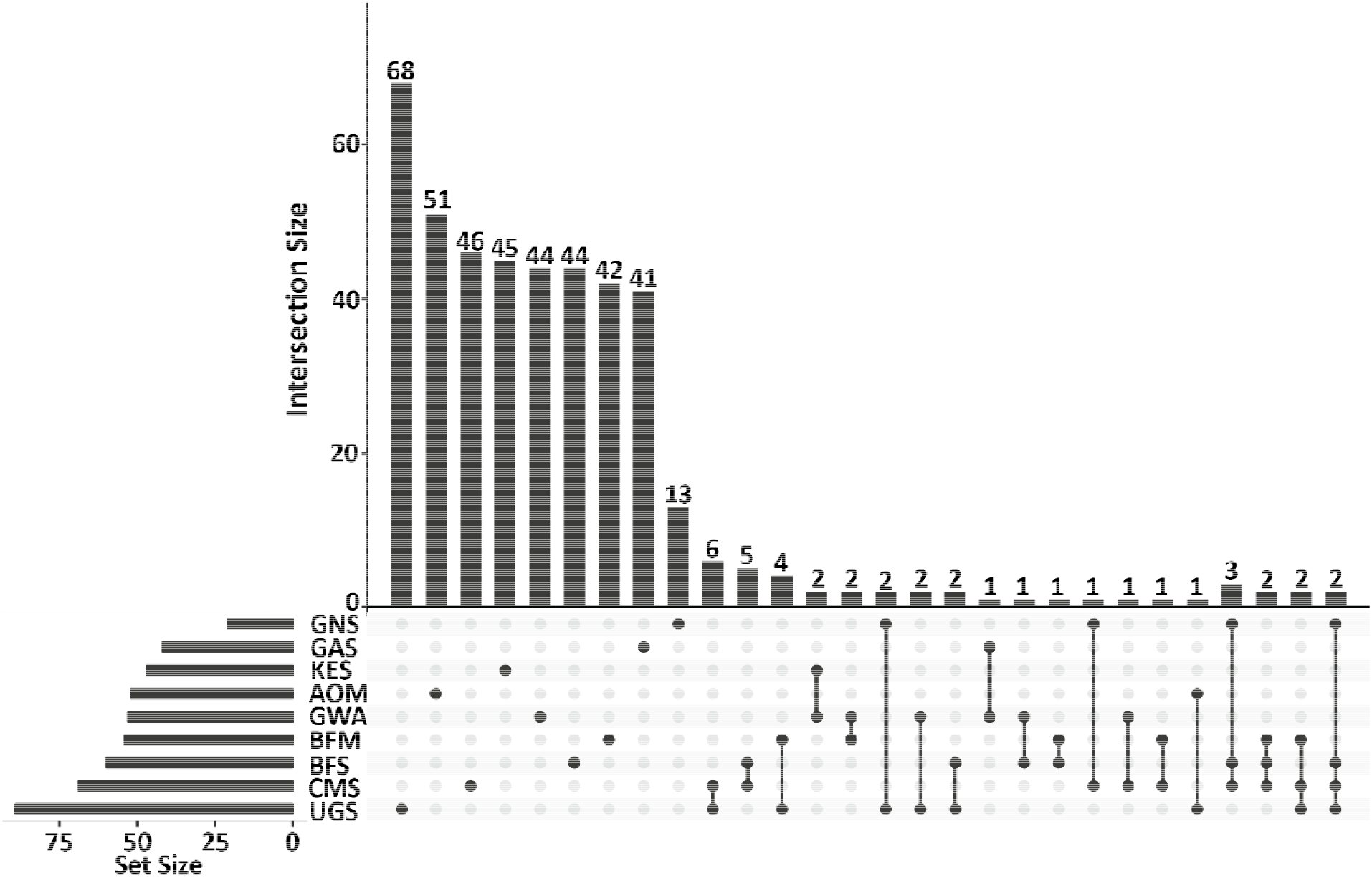
An Upset analysis depicting the distribution of unique and shared recombination hotspots. Within this figure the set size represents the total number of hotspots found within each population, while the intersection size defines the number of hotspts that were identified in a single population (single dots) and the number of hotspots shared across multiple populations (connected dots). The numbers at the top of each bar define the number of hotspost found within that interesection.

Despite finding very little evidence of overlapping (shared) hotspots, empirical overlaps were on average larger than expected when hotspots were randomly shuffled throughout the genome, resulting in a positive net difference (empirical hotspot overlap – shuffled overlap; Supplementary Table S6). Furthermore, pairwise Fishers Exact test showed that a total of 19 (out of 26) comparisons had significantly larger hotspot overlaps than stochastic expectation (p < 0.05; Supplementary Table S6). Of the 19 significant comparisons, most were found between populations of *A. gambiae* (14 comparisons) while the remaining comparisons were between *A. gambiae* and *A. coluzzii*; there were no significant hotspot overlaps within *A. coluzzii*.

### Consequences of recombination hotspots on population diversity, and divergence

Average nucleotide diversity was larger within recombination hotspots (0.02578 ± 0.00073) compared to genomic background estimates (0.02257 ± 0.00002201) (Figure 6). Across the genome, we show that within most populations, nucleotide diversity was significantly larger (p < 0.05) within recombination hotspots than compared to the surrounding background levels (Figure 6). Moreover, for autosomal chromosomes, the average nucleotide diversity was significantly larger within recombination hotspots (0.02893 ± 0.00038) than compared to the average background (0.02567 ± 0.00001322). Across the X chromosome, nucleotide diversity followed similar patterns where nucleotide diversity was significantly larger within recombination hotspots (0.02264 ± 0.00109) than compared to the genomic background (0.01947 ± 0.0000308) (Figure 6).

**Figure 6.**
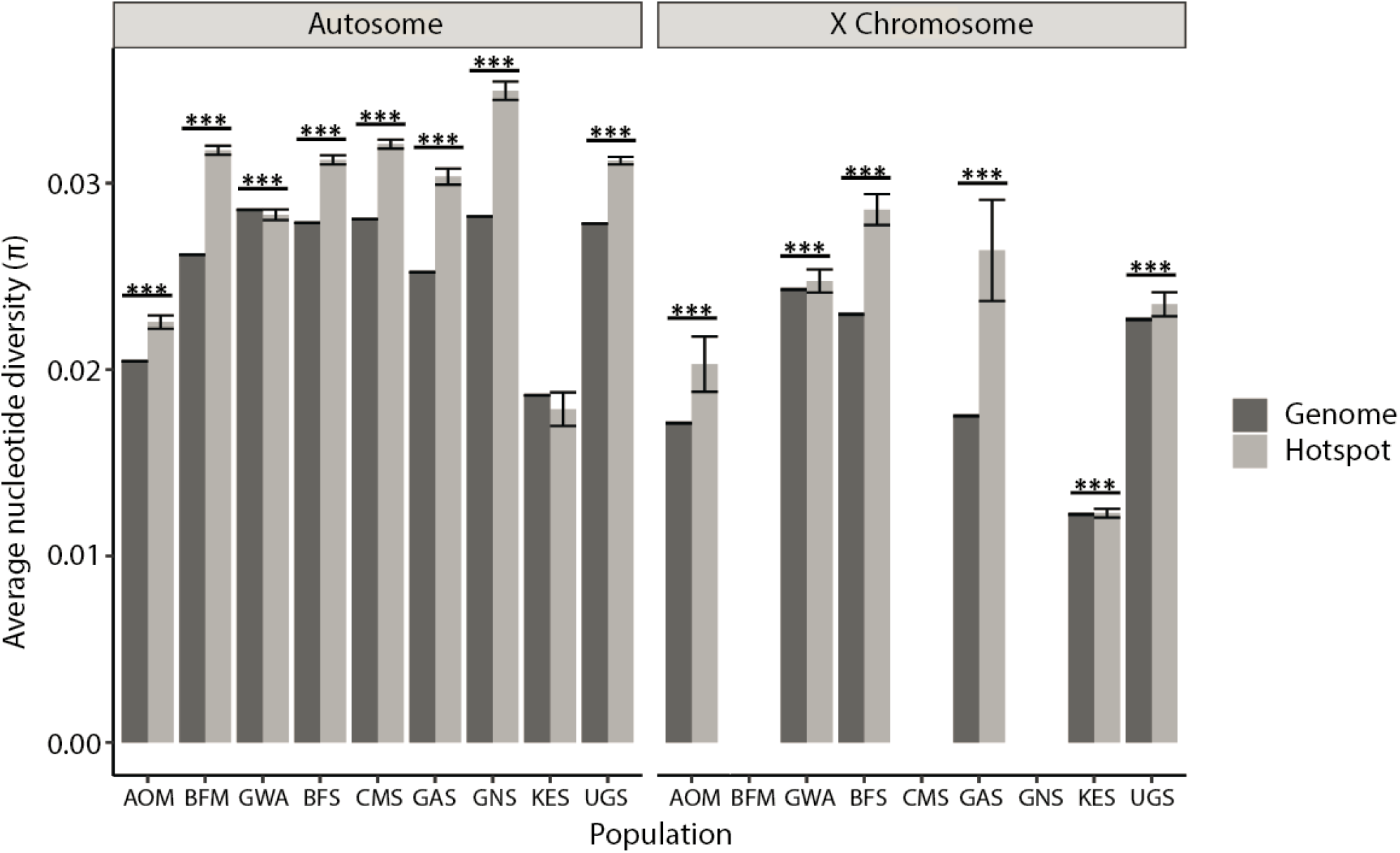
Barplot comparing nucleotide diversity within recombination hotspots and genome-wide nucleotide diversity for the autosomomes and X chromosomes. Vertical lines at the end of each bar represents the standard error for each measurement. Solid horizontal bars and asterisks denote significant differences between recombination hotspots and the overall genome. The asterisks above the bars denotes the level of significance (* = p-value ≤ 0.05, ** = p-value ≤ 0.01, and *** = p-value ≤ 0.001).

Similar to nucleotide diversity, average estimates of dxy proved to be larger within recombination hotspots (0.0244 ± 0.0004) when compared to a genome-wide estimate (0.0215 ± 0.0000122). Between autosomal and the X chromosome most population pairwise comparisons support evidence suggesting significant differences (p < 0.05) in the number of per site differences within recombination hotspots (Figure 7A). Moreover, across autosomal chromosomes average dxy was larger within recombination hotspots (0.0272 ± 0.000325) than average dxy estimates across the genome (0.0241 ± 0.0000113). For the X chromosome, average dxy estimates were larger within recombination hotspots (0.0215 ± 0.000475) than compared to dxy along the X chromosome (0.0189 ± 0.0000130) (Supplementary Figure S10). Regions within recombination hotspots experience ~3.0% more nucleotide differences per site than the entire X chromosome.

**Figure 7.**
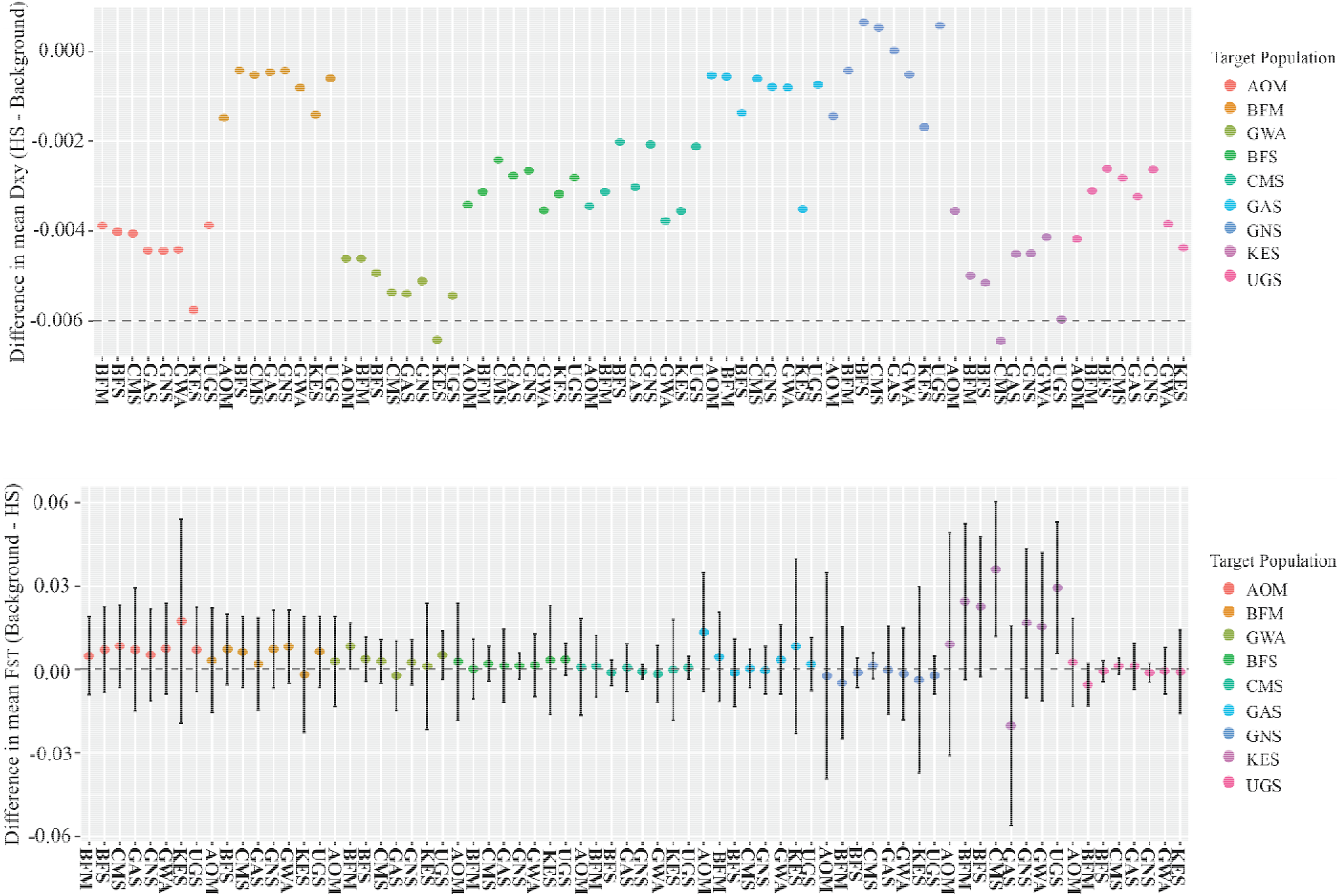
Barplots that show the ratio between dxy (top panel) and F_ST_ (bottom panel) within recombintion hotspots compared to background estimates for autosomal chromosomes. For each plot, the target population is the population in which within recombinaition hotspts are being compared to the same genomic regions withi the populations labeled along the X-axis.. The dashed black line signifies no difference in divergence metrics (dxy or F_ST_) between the genomic background and hotspots. The astrixes above represents significant pairwise comparisons.

When we examine the patterns of F_ST_ we find that of the 72 comparisons, 51 (70.8%) showed an increase in F_ST_ within recombination hotspots, whereas the remaining 21 comparisons (29.2%) showed a decrease in F_ST_ within hotspots (Figure 7B). However, the difference ratio between the genomic background F_ST_ and adjacent hotspot F_ST_ zero, suggesting very little change in relative genetic differentiation between hotspots and the adjacent background for all comparisons. Despite this, we still identified 13 comparisons where F_ST_ was significantly greater than what was predicted from the distribution of F_ST_ for non-hotspot regions. Variance of F_ST_ followed similar patterns where across all hotspots it was higher (0.00742) when compared to the F_ST_ variance of the adjacent genomic backgrounds (0.00702); however, this difference was not significant (p = 0.0660).

## Discussion

In this study, we report the first recombination landscape for multiple populations of *A. gambiae* and *A. coluzzii*. We identified large variations in recombination rates across all populations. We showed that effective population size, chromosomal location (centromeric regions), chromosomal inversions, and selective sweeps all contribute to recombination differentiation among populations. We identified several hotspots on both X and autosomal chromosomes suggesting regions of the genome with significantly higher recombination rates. Lastly, we showed significant associations between recombination hotspots, nucleotide diversity, and relative/absolute divergence, suggesting that recombination hotspots may be playing a role in not only maintaining within population diversity, but also contributing to between population divergence.

### Fine-scale recombination rates across populations of Anopheles

Consistent with other study systems, our results suggest differences in recombination rates among populations (Kong et al. 2002; Nachman 2002; Smukowski and Noor 2011) (Figure 2; Table 1). Our results suggest that these differences are not just the consequence of overall changes in recombination rate among populations, but the result of differences in fine-scale recombination rates along the genome (Table 2), consistent with recent findings (Spence and Song 2019). Importantly, these patterns were consistent with previous studies showing high levels of recombination divergence at fine scale, whereas more conservation in recombination rates across the entire genome (Smukowski and Noor 2011). It has been proposed that variation in recombination rates is due to evolutionary constraints that allows for a lower and upper bound of recombination (Smukowski and Noor 2011; Ritz et al. 2017). Lower recombination rates may prove to be maladaptive as the genome lacks the ability to create novel haplotypes and reduce interference of deleterious mutations. Conversely, extremely high levels of recombination have the potential to break up beneficial haplotypes and gene complexes, proving to be maladaptive. These bounds may be responsible for the conservation of the average genome-wide recombination rates, however, the localized differences in recombination may represent the variation in recombination within evolutionary constraints., or the presence of selective sweeps. Regions of the genome with recently fixed selective sweeps might display excessive LD and low recombination (Kim and Nielsen 2004; Sibley and Ajioka 2008). We see this pattern reflected in the recombination landscape of *Anopheles* when we focus on genes involved in insecticide resistance, and more thoroughly discuss this results in the following sections.

### Evolutionary and genomic features shaping the recombination landscape

We showed that the recombination landscape is heavily influenced by evolutionary and genomic features such as effective population size, chromosomal location, inversions, and selective sweeps. For example, populations that are south of the Congo River basin (AOM and GAS) and east of the African Rift valley (KES) have shown genomic evidence of recent bottlenecks (Lehmann et al. 1999; Anopheles gambiae 1000 Genomes Consortium 2017). Indeed AOM, GAS, and KES had the lowest variance, as well as reduced population-scaled recombination (ρ/kb) (Figure 2A). However, when calculating the per site recombination rate to account for effective population size, KES had an average recombination rate that was an order of magnitude larger than the other populations. As demonstrated within previous studies, populations with a small effective population size may exhibit higher recombination rates associated to an increase accumulation of deleterious mutations in homogenized genomes (Keightley and Otto 2006; Kumar et al. 2019; Schwarzkopf et al. 2020). Because KES has had a small historical Ne (~10,000), we hypothesize that the deleterious effects of a low Ne have contributed to a increased overall recombination rates within KES; however, this hypothesis needs to be tested.

Chromosomal location is also a major influence on the recombination landscape. For example, recombination was reduced near the centromeric regions for all populations (Figure 2 & 3). Reduced recombination near centromeres has been previously observed (Nambiar and Smith 2016). Recombination within centromeric regions is often considered maladaptive where it can break up conserved blocks of genes and interfere with proper chromosome segregation during meiosis, which has the potential to create aneuploid progeny (Nambiar and Smith 2016). In the case of *A. gambiae* and *A. coluzzii*, each species is grouped by a molecular form based off diagnostic genetic differences in the ribosomal DNA, near centromeric regions of the X chromosome (Coetzee et al. 2013; Aboagye-Antwi et al. 2015). These fixed differences have been shown to play a role in positive assortative mating, thus reducing gene flow between the two species (Aboagye-Antwi et al. 2015). It has long been speculated that low levels of recombination rates along the X chromosome have contributed to reproductive isolation between *A. gambiae* and *A. coluzzii*. Here, for the first time, we show drastic reduction in recombination rates along the centromeric regions of the X chromosomes across all sampled populations that supports the underlining hypothesis of little recombination near the centromere of the X chromosome.

Of the six inversions within *Anopheles*, two of them, 2Rb and 2La, show prominent associations for specific environments (rain-dependent vs. rain-independent) (Coluzzi et al. 1979; Lanzaro and Lee 2013; Cheng et al. 2018). For populations that are homozygous for either chromosomal inversion (2La or 2Rb), or the wild type (no inversion), there were no reductions in recombination rates across the 2La and 2Rb chromosomal inversions. Indeed, the inversion loop does not need to form during meiosis (Figure 3; Table 3), and therefore, does not result in maladaptive indels within recombinant chromosomes. However, for populations that were polymorphic for the inversions, recombination rates within the inversion were largely suppressed. This drastic reduction in recombination was also followed by an increase in recombination along regions that flank the inversions (Figure 3; Table 3). Though, for populations that were polymorphic for chromosomal inversions we still detected evidence for successful recombination events within these inversions, and in some cases, recombination hotspots. Successful recombination within inversions could be explained by a double cross-over within inversion loops which, can potentially prevent the major insertions/deletions that occur between inverted and non-inverted homologous chromosomes, resulting in successful recombinant progeny. Previous studies focusing on the clinal variation of chromosomal inversions have suggested that relaxed pressure for the 2La inversion in non-arid regions may lead to an increase in recombination (through double cross-overs) within inverted regions (Cheng et al. 2012).

### Recombination rates within insecticide resistance genes

Because of long-term exposure to insecticides, recent selection analyses show signatures of molecular adaptation across several genes known to be involved with insecticide resistance (Anopheles gambiae 1000 Genomes Consortium 2017). Although this exposure has been relatively long in an ecological timeframe; the selection imposed by insecticides on *Anopheles* populations have only been acting recently in an evolutionary time scale. The implications for the evolution of resistance in these populations is that we expect to see reduced recombination in areas surrounding the genes involved in insecticide resistance. These regions are of great importance because as insecticide resistance increases, *Anopheles* becomes increasingly difficult to control and indirectly making the spread of malaria tough to mitigate. Recombination is especially important because of its ability to generate novel haplotypes which may increase insecticide resistance, especially under models that apply an evolutionary arms race (Clay and Kover 1996). However, here we show that within IR genes, especially within Cyp6p and Cyp9k1 (both known for the resistance towards DDT, and pyrethroid-based compounds), recombination is reduced within 1-3Mb of the IR gene (Figure 4; Supplementary Figure S2-S7). These genes were found to be under selection in UGS and BFM, however, they are neutrally (or show a weak signal of selection) evolving in other populations such as GAS and AOM (Anopheles gambiae 1000 Genomes Consortium 2017). For those populations with neutrally evolving IR genes, we found no change in the recombination rates within the same regions (See Figure 4). For those populations with neutrally evolving IR genes, we found no change in the recombination rates within the same regions. This is consistent with the recent selective sweeps containing little variation and reduced signature of recombination. An alternative explanation, is that recombination within IR genes may break apart favorable haplotypes and otherwise prove to be maladaptive in terms of insecticide resistance, with natural selection preventing the spread of recombinants in the population. Perhaps a comprehensive study involving the differences in fitness across recombinant and parental chromosomes when exposed to varying levels of insecticides could help us shed some light between the role that recombination and selection play in the evolution of this locus.

### Identifying recombination hotspots

Because of the absence of recombination hotspots in *Drosophila* (Chan et al. 2012), a main question in recombination studies is if this pattern is true across other insect systems, including those that are closely related to *Drosophila*? Implementing LD methods, which have not been applied to *Drosophila* systems in order to identify hotspots, we identified a total of 436 genomic regions that were consistent with recombination hotspots within *Anopheles* (Figure 5; Supplementary Table S5). Majority of recombination hotspots were unique to a specific population, however, only 9.6% (40 hotspots) were shared between two or more populations.

For all population pairwise comparisons that shared a hotspot, all but one had empirical overlaps that were larger than the overlaps generate from stochastic hotspot shuffling (Supplementary Table S6). Moreover, for the 25 comparisons where the empirical overlap was larger than the simulated overlaps, Fisher’s Exact test showed that 19 comparisons were statistically significant (p-value < 0.05), suggesting a sharing in recombination hotspots more so than what is expected under a stochastic assumption (Supplementary Table S6). It should be noted that most of the significant overlaps were within populations of *A. gambiae*, however there were few significant overlaps between *A. gambiae* and *A. coluzzii*, and none within *A. coluzzii*. This is a very interesting finding because it demonstrates the potential conservation of few recombination hotspots within the *Anopheles* genome, which may be serving a functionally important role in the generation of novel variation within the genome.

### Consequences of recombination hotspots on population diversity, and divergence and implications for the identification of adaptive variation

Within autosomal chromosomes, we identified significant differences between nucleotide diversity within recombination hotspots and average genomic levels. Specifically, nucleotide diversity was significantly higher in recombination hotspots for all populations with the exception of KES (Figure 6). Because of the recent, and severe, bottleneck for *Anopheles* within Kenya, it is not surprising that there were no significant differences in nucleotide diversity. Similarly, for the X chromosome, all nucleotide diversity comparisons also proved to be significant (Figure 6). Identifying significant increase in nucleotide diversity within recombination hotspot is an interesting finding because it suggests an evolutionary role for recombination in the maintenance of genetic variation. More specifically, the presence of recombination hotspots could be acting as protected reservoirs for genetic variation by reducing the effects of linked selection.

Much like nucleotide diversity, the number of per-site differences (dxy) between populations followed similar patterns along the autosomal chromosomes, where dxy within recombination hotpots were significantly larger than background estimates for 66 of 72 (91.6%) comparisons, suggesting an increase in population divergence within these regions (Figure 7A). The only population comparisons that did not show a significant increase in dxy within hotspots were within AOM and KES. These two populations have smaller effective population sizes and could be a contributing factor to why these populations experienced lower overall changes in dxy. Similarly, along the X chromosome, we identified significant increases in dxy within recombination hotspots for 43 of 46 (93.4%) comparisons, suggesting that population divergence is significantly larger within these regions (Supplementary Figure S10). Much like the autosomes, two comparisons within AOM and one within GAS did not reflect significant differences in dxy. Effective population size for GAS was estimated to be on the same order of magnitude as AOM, further supporting an increased role of genetic drift across the genome resulting in higher background levels of genetic divergence. With the exception to nine comparisons, the remaining comparisons support a mutagenic hypothesis where the mechanistic properties of recombination not only influence variation within a population, but also drives localized regions of the genome to have elevated levels of divergence. The other nine comparisons support a linked selection hypothesis where recombination is maintaining genetic diversity within populations, but elevated levels of divergence are larger where recombination rates are lower. Importantly, these results suggest that recombination hotspots are maintaining, and potentially generating, genetic diversity and contributing to absolute genetic divergence within and between species of *Anopheles*.

The relationship between the recombination landscape and F_ST_ is more complex than patterns of diversity and divergence (as measured by dxy). For example, we showed that only 29.2% of our pairwise comparisons suggest that F_ST_ was lower within recombination hotspots when compared to the average genome (Figure 7B). Despite the differences in background and hotspot genetic differentiation being close to zero (suggesting little difference between the two comparisons) 13 pairwise comparisons were significantly higher F_ST_ within recombination hotspot compared to background estimates. These empirical data suggest that elevated recombination rates may not play an essential role in reducing levels of genetic differentiation. However, our results provide little insight on the efficacy of elevated F_ST_ within regions of low recombination and warrants further investigation. Patterns of genetic differentiation are a common statistic in population genetics used to infer differences in allele frequencies and sometime selection. Here we provide evidence that suggests that regions of the genome with increased recombination rates can retain genetic variation and suppress patterns of genetic differentiation between populations. However, the data also suggest that elevated recombination rates can still lead to patterns of increased genetic differentiation. Overall, we find that the recombination landscape can have significant impacts on patterns of genetic diversity, divergence, but potentially a smaller impact on patterns of F_ST_. We suggest that future studies involved with identifying patterns of genetic diversity and divergence across populations also incorporate the recombination landscape, to better infer the patterns of genetic variation.

## Conclusion

In this study, we detailed the fine-scale recombination maps and hotspot locations across nine populations containing two species of *Anopheles*; *A. gambiae* and *A. coluzzii*. We identified patterns of recombination rates that are similar to other systems that show large rate variation. We also show that changes in the landscape of recombination maps strongly depends on location along the chromosome, chromosomal inversions, and selective sweeps within IR genes. The changes in recombination rates along the genome are even more prominent near the centromeric regions of the X chromosome where recombination rates are minimal and may be responsible for maintaining species boundaries between *A. gambiae* and *A. coluzzii*. The proportion of unique and shared hotspots support the findings of other studies suggesting that there is a quick evolutionary turnover of hotspots and recombination rates; however, few hotspot locations may be more conserved, potentially serving important biological functions within the genome. Lastly, we show evidence that recombination hotspots have the potential to increase both nucleotide diversity within populations and the number of nucleotide differences between populations. This was also true for patterns of relative genetic differentiation; however, these results were less consistent and show that elevated recombination can also reduce levels of genetic differentiation. Overall, our results provide significant insight into the evolutionary role of recombination across populations of *Anopheles*, which can be implemented for further understanding evolutionary trajectory of a prominent biological vector.

## Materials and Methods

### Sample and Sequence data

For this study we used full genome data from previously published work that was aimed at characterizing the genomic structure, patterns of selection, migration, and estimates of effective population size (*N_e_*) for *A. gambiae* and *A. coluzzii* (Anopheles gambiae 1000 Genomes Consortium 2017). Specifically, sequencing was performed on an Illumina HiSeq 2000 platform at the Wellcome Trust Sanger Institute. All sequence reads were aligned to the AgamP3 reference genome (Sharakhova et al. 2007) using bwa (Li and Durbin 2009) and Single Nucleotide Polymorphisms (SNPs) were discovered using GATK under their best practices protocol (DePristo et al. 2011; Van der Auwera, Geraldine A et al. 2013). For more information pertaining to the sequence data used and sample collection see (Anopheles gambiae 1000 Genomes Consortium 2017). We used a total of 441 individuals (882 genomes) sampled over nine populations, eight countries, and two species of *Anopheles*. Sample sizes ranged from 88 to 120 genomes per population (Figure 1; Supplementary Table S7). Prior to estimating recombination rates, we filtered bi-allelic phased SNP data to account for a minor allele frequency of at least three individuals, missing data, and fixed sites.

### Estimating fine-scale recombination rates

To estimate fine-scale recombination rates, we used LDhat which is a package of programs that estimates population recombination rates by implementing the composite likelihood method of Hudson et al. (2001) (Hudson 2001; McVean, Gil and Auton 2007). Because obtaining fine-scale estimates of recombination across a genome for a large number of samples is computationally exhaustive, we decomposed all genomic data in two different components to help reduce computational resources. First, LDhat was ran in parallel within each population allowing the split of individuals and reduced memory usage. Second, instead of estimating recombination rates for the entire genome at one time, each chromosome arm was divided into sliding windows, further reducing the amount of memory required for each population/chromosome. Each window contained 2,000 SNPs including a 500 SNP overlap between adjacent windows and ran for 100,000,000 iterations with a burn-in period of 50,000,000 iterations. LDhat sampled every 10,000 iterations, providing 10,000 sampled recombination rates per site under the INTERVAL function. After obtaining recombination rates the STATS function was implemented in LDhat which calculates the mean, median, upper 95%, and lower 95% recombination rate for each SNP. Windows were then trimmed of their overlaps and aligned to their respected positions along the chromosome. This pipeline was performed on both autosomal and X chromosomes, however, within the *Anopheles* system males are the heterogametic sex. We, therefore, included only females for building the recombination maps for the X chromosome. In total we used 417 females for estimating recombination rates along the X chromosome (see Supplementary Table S7).

### Detecting recombination hotspots

In order to detect recombination hotspots, we used LDhot, which is a program designed to identify regions of the genome where recombination rates are significantly larger than background levels (McVean, G. A. et al. 2004; Myers et al. 2005). Specifically, LDhot defines a Poisson distribution with a given lambda coupled with a sliding window approach to identify regions of the genome that have recombination rates explained by a different lambda and Poisson distribution (McVean, G. A. et al. 2004). If localized patterns of recombination rates are better explained by a different Poisson distribution, then LDhot flags the window as a recombination hotspot. Poisson distributions were defined based off 1,000 simulations. Only sliding windows with a p-value < 0.01 were retained and then merged together using BedTools MERGE (Quinlan and Hall 2010) to identify the size and location of the recombination hotspot.

However, because LDhot is known to lose power when population-scaled recombination rates are low (ρ < 5) (Johnston and Cutler 2012; Wall and Stevison 2016) we retained hotspots that only showed large deviations in recombination rates from the surrounding background. Specifically, we compared recombination rates within hotspots to the recombination rates for the surrounding 50 kb background (the same size flanks that LDhot uses to detect hotspots). We used the difference in recombination rates between hotspots and the adjacent 50kb background to rank the intensity of recombination hotspots along a given background (supplemental Figures S11-S16). We retained only the top 5% of hotspots with the largest difference in rho when compared to the surrounding background rates for each population. Though this method is conserved, we have a greater confidence in our filtered hotspots as being true hotspots and not an artifact of demographic and computational limitations.

After hotspots were identified, we used UpSet in R to view the union and intersection of hotspots for all population pairwise comparisons (Conway et al. 2017). For all pairwise comparisons that shared at least one recombination hotspot, we wanted to test if these hotspots were more likely shared due to stochastic processes (random hotspot location within the genome) or if there was potential evidence of hotspot conservation between populations (shared more than random expectations). Here, the degree of overlap was determined by how many base pairs were shared within a given recombination hotspot. Specifically, to perform stochastic shuffling of recombination hotspots, we used BedTools SHUFFLE (Quinlan and Hall 2010) to randomly shuffle known hotspot locations for each chromosome arm. Recombination hotspots were shuffled for 10,000 iterations where the degree of overlap (in bp) was calculated for each iteration. The differences between average simulated and empirical overlaps were then calculated; a positive net difference suggests that empirical overlaps were larger than the shuffled overlaps. To complement stochastic simulations, we performed a Fishers Exact Test, using BedTools, to identify significant deviations from our null hypothesis of stochastic hotspot sharing across all empirical overlaps. We defined a significant overlap with a p-value < 0.05.

### Consequences of elevated recombination on diversity and divergence

We compared levels of diversity within and between populations to understand potential consequences that recombination has on genetic diversity and divergence. For this, levels of nucleotide diversity (π) were calculated within recombination hotspots and compared to the per site genome-wide estimates of π for each population. Because of prominent reproductive isolation along the X chromosome, this analysis was performed on the autosomal and X chromosome separately. To determine significant differences for π between hotspots and the genomic background a Wilcoxon Rank test was performed; a non-parametric statistical test that does not assume the data are normally distributed, which is the case for estimates of π (p-value < 0.05). To test for elevated levels of divergence within recombination hotspots we calculated two different metrics of genetic divergence: absolute divergence (dxy) and relative divergence (F_ST_). Specifically, for dxy, we calculated the number of per site differences for each recombination hotspot for each population pairwise comparison and compared dxy estimates across the average genomic background level.. A Wilcoxon Rank Test was performed for each dxy pairwise comparison was used to identify significant deviations (p-value ≤ 0.05).

We used a randomization approximation to test the significance of the difference in F_ST_ between regions with HS of recombination and Background. For this, each the F_ST_ values for each region were labeled as HS or Background, and a shuffling procedure of the labels was used to generate 1000 pseudo-replicates. The average difference in F_ST_ between relabeled HS and Background regions was estimated for each pseudo-replicate and a distribution was generated. An empirical p-value (p-value ≤ 0.05) for the significance of the observed difference was determined by looking at the proportion of pseudo-replicates with values larger than 95% of the observed difference.

## Notes

https://www.malariagen.net/data/ag1000g-phase1-ar3.1

### Competing Interest Statement

The authors have declared no competing interest.

